# Siglec-F Deficiency Prevents Fibrosis After Bleomycin-Induced Acute Lung Injury

**DOI:** 10.1101/2024.10.04.616703

**Authors:** Marika Orlov, Sunad Rangarajan, Ana M. Jaramillo, Qihua Ye, Naoko Hara, Kenny Ngo, James C. NeeDell, Anna Q. Harder, Fan Jia, Brian Vestal, Rachel Z. Blumhagen, Ting-Hui Tu, Jazalle McClendon, Alexandra L. McCubbrey, Bradford J. Smith, David A. Schwartz, William J. Janssen, Christopher M. Evans

## Abstract

Injury to the lungs causes acute inflammation that can lead to pathological lung fibrosis. Airspace macrophages (AMs) are critical for repair of injured tissue, but they can contribute fibrosis through mechanisms that are incompletely understood. Siglecs are expressed by immune cells. In mice, Siglec-F is chiefly expressed AMs where it is considered inflammosuppressive. We hypothesized that its deletion would worsen lung injury and fibrosis in response to intratracheal bleomycin challenge. We evaluated Siglec-F expression and function in mice challenged with bleomycin on days 7, 14, and 21 post-challenge (2.5 U/kg). AMs were the predominant inflammatory cells at all timepoints, and they included resident (RAM) and recruited (RecAM) subsets. Siglec-F deficiency prevented fibrosis than in Siglecf−/− mouse lungs, as evident from biochemical and histologic readouts. We performed RNAseq on pooled RAMs and RecAMs from wild type and Siglec-F deficient mice. Lung fibrosis 21 d after bleomycin challenge was associated with differentially expressed genes (DEGs) related to cholesterol synthesis and metabolism. In AMs from healthy lung lavage fluid and idiopathic pulmonary fibrosis patient tissues, the human paralogs Siglec-7 and Siglec-9 were expressed. Findings here identify novel mechanisms that control protective and detrimental functions of AMs after lung injury.

---

Injury to the lungs causes acute inflammation followed by resolution and tissue repair. In some patients who recover from acute lung injury, excessive scarring can lead to fibrosis that is persistent and is presently irreversible (1). To identify potential treatments for lung fibrosis, there is a need to determine causative mechanisms and the cell types involved. Airspace macrophages (AMs) are critical for resolving inflammation and facilitating repair of injured lung tissue, but they can also contribute to the development of fibrosis thereby implicating AMs as putative targets (2, 3).

After injury, AM pools expand markedly through local replication of resident airspace macrophages (RAMs) and through entrance of circulating monocytes into the lungs where they mature into recruited AMs (RecAMs) (4). RAMs and RecAMs are typically distinguished by surface markers including CD11c (RAMs) and CD11b (RecAMs). They can be further segregated in mouse models using the surface protein Siglec-F whose expression is high in RAMs and low in RecAMs (2). Despite its common use as a marker, Siglec-F function in AMs is unknown.

Siglecs are sialic-acid-binding immunoglobulin-like lectins that are widely expressed by immune cells and comprise numerous inhibitory immunoreceptors (5, 6). In mice, Siglec-F is chiefly expressed by AMs and eosinophils (5, 6). In eosinophils, Siglec-F and its human paralog Siglec-8 inhibit allergic inflammation (7, 8). Since Siglec-F is considered inflammosuppressive, we hypothesized that its deletion could worsen injury in the lungs, thereby exacerbating fibrosis. To address this, we evaluated Siglec-F expression and function in mice challenged with bleomycin (2.5 units/kg, IT) or saline vehicle. Inflammation and fibrosis endpoints were collected 7, 14, and 21 days post-exposure. Detailed methods are in the data supplement.

In lung lavage fluid, AMs were the predominant inflammatory cells found at all timepoints. AM numbers peaked on day 7 (Figure 1A), consistent with migration of RecAMs into the lungs (2). Initially, RecAMs did not express Siglec-F, but within 14 days, Siglec-F increased to ∼75% of the levels observed in RAMs (Figure 1B). These data supported our overarching hypothesis that AM responses to bleomycin-induced injury involve Siglec-F. To determine its significance, we tested the effects of Siglec-F deficiency on fibrotic responses to bleomycin.

**Figure 1.**
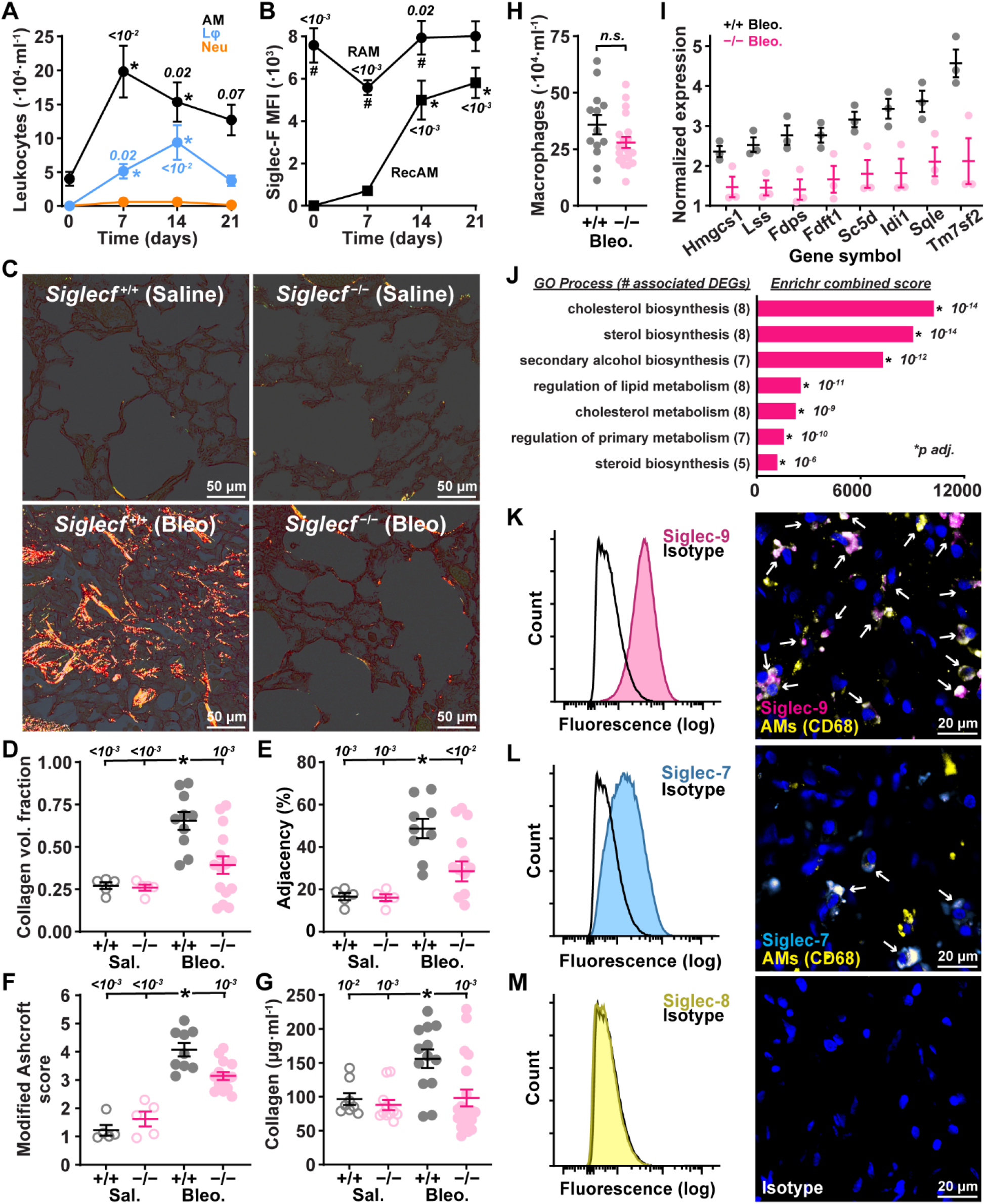
Siglec expression and function by airspace macrophages in lung fibrosis. **A**. Airspace macrophage (AM), lymphocyte (Lφ), and neutrophil (Neu) populations were quantified in bronchoalveolar lavage (BAL) fluid at baseline (n=4) and on days 7 (n=8), 14 (n=8), or 21 (n=6) after bleomycin challenge (bleo, 2.5 U/kg, IT). **B**. Resident (RAM, CD11b-, Circles) and recruited (RecAM, CD11b+, Squares) AM subsets were evaluated for Siglec-F expression by flow cytometry over the same 0-21 day timecourse (n=6, 7, 11, and 9, respectively). **C-G**. Fibrosis was assessed in lungs from wild type (+/+) and *Siglecf*^-/-^ (-/-) mice 21 days after bleo or saline vehicle challenge. Picrosirius red staining showed collagen birefringence under polarized light (**C**). Stereology was used to quantify collagen volume fraction (**D**), and the adjacency of positive points was used to estimate collagen density (**E**). Findings were validated using Ashcroft score (**F**) and lung hydroxyproline content (**G**) measurements. **H-J**. To assess AM programming, BAL AMs were quantified (**H**), purified by FACS, and analyzed by RNAseq. DESeq2, limma, and Enrichr identified differentially expressed genes (DEGs) related to lipid or cholesterol metabolism that were significantly reduced in bleo challenged *Siglecf*^-/-^ mice compared to bleo challenged wild type controls (**I**,**J**). **K-M** Comparative analysis of human siglec paralogs was performed with flow cytometry using AMs isolated from BAL of healthy humans and immunofluorescence using IPF tissues. Human AMs express Siglec-9 (**K**) and Siglec-7 (**L**). Siglec-8 (**M, left**) was not detected in healthy AMs, and isotype control labels showed no signal in IPF lungs (**M, right**). Black lines in flow cytometry represent IgG controls for each antibody in **K-M**. Significance was determined using ANOVA with multiple comparisons. Data were analyzed by Kruskal-Wallis test (‘*’, sig. vs. day 0; # sig. between RAM and RecAM) in **A** and **B;** ANOVA (‘*’, sig. vs. bleomycin-challenged wild type), and two-way Mann-Whitney test in **H**. In **J**, adjusted p-value (p-adj.) is the z-score of deviation from the expected rank by the Fisher exact test. Significance cut-off was 0.05 for all data.

Counter to our initial hypothesis that Siglec-F deficiency would worsen fibrosis, we observed less fibrosis than in *Siglecf*^*−/−*^ lungs than wild type. Picrosirius red staining showed reduced collagen on day 21 post-bleomycin (Figure 1C). When quantified using a blinded stereologic analysis of collagen fiber birefringence under polarized light (Figure E1), the volume fraction of collagen and its distribution were significantly lower in *Siglecf*^*−/−*^ lungs (Figure 1D,E). For further validation, we quantified fibrosis using modified Ashcroft scoring and hydroxyproline assays (9). Both validated our observation that bleomycin-induced lung fibrosis was suppressed in *Siglecf*^*−/−*^ mice (Figure 1F,G).

The results above provided strong evidence for pro-fibrotic processes that were dependent on Siglec-F expression in AMs. Although, AM numbers in BAL were not significantly different in bleomycin challenged wild type and *Siglecf*^*−/−*^ mice (Figure 1H, Table E1), we postulated that there could be Siglec-F-dependent differences in AM programming. We performed bulk RNAseq on AMs collected from wild type and Siglec-F deficient mice. Given the prominence of Siglec-F as an AM marker, its absence reduced our ability to distinguish RAMs from RecAMs. This was further hindered by findings that RecAMs acquire CD11c over the timecourse used here (10). Hence, we conducted RNAseq on total AM populations containing both RAM and RecAM subsets.

On day 21 after bleomycin challenge, lung fibrosis was associated with 24 differentially expressed genes (DEGs) in AMs (Figure E3). Enrichr pathway analysis revealed significant overlap with cholesterol synthesis and metabolism (11). This was attributed to 8 DEGs--*Lss, Idi1, Sc5d, Hmgsc1, Tm7sf2, Fdps, Fdft1*, and *Sqle*--all of which were down-regulated in *Siglecf*^*−/−*^ mice (Figure 1I,J). While these data did not reveal a canonical pro-fibrotic signature in AMs, they did identify putative pathways that may be involved in tissue repair. Future analyses using tools that better separate RAMs and RecAMs may help further refine these.

Human siglecs have not been studied extensively in the context of acute lung injury leading to post-inflammatory fibrosis. Our prior bulk and single-cell RNA sequencing on RAMs isolated from healthy human lungs showed moderate to high expression of the human paralogs of Siglec-F: Siglec-7, -8, and -9 (3, 12). To verify transcriptomic results, we used flow cytometry to examine Siglec-7, -8, and -9 proteins on AMs in bronchoalveolar lavage fluid from healthy human lungs and immunofluorescence to evaluate siglec expression in human IPF tissues.

Siglec-9 and Siglec-7 were both expressed in healthy AMs, and they were also observed in AMs from patients with pulmonary fibrosis (Figure 1K,L). By contrast, Siglec-8 expression was absent (Figure 1M). These results demonstrate that Siglec-9 and Siglec-7 are expressed on human AMs in health and in pulmonary fibrosis where they may mediate inflammation and repair functions similar to those mediated by Siglec-F in mice.

In summary, these studies reveal the potential importance of Siglec-F signaling in AMs during resolution of acute lung injury. Siglec-F is important for lung repair as *Siglecf*^*−/−*^ mice are protected from fibrosis following acute injury. Bleomycin challenge triggers periods of injury, fibroproliferation, and fibrosis that are coupled with inflammatory, resolving, and reparative AM programming phases (2, 9). It has been postulated that macrophages utilize shifts in cholesterol metabolism to alter their responses inflammation and tissue damage (13). The exact role for shifts in AM cholesterol metabolism and the development of fibrosis are incompletely understood, but aberrant cholesterol metabolism is linked to macrophage-mediated fibrotic responses in mice (14) and potentially to IPF pathogenesis in humans (15). Our results provide a possible mechanistic link to AM effector functions, and additional studies are needed to identify endogenous Siglec-F ligands such as sialylated Muc5b (8) and to delineate selective contributions of AM subsets in Siglec-F dependent inflammation and fibrosis (2). Findings here lay the foundation for further experiments focused on mechanisms that control protective and detrimental functions of AMs in lung injury and fibrosis.

## Acknowledgement of grant support

R01HL080396, I01BX005343 (CME); R01HL14974, R35HL140039 (WJJ); R01HL130938 (CME,SR,WJJ); 5T32HL007085 (MO); R01HL158668, R01HL149836, I01BX005295, UG3/UH3HL151865 (DAS); P01HL162607 (DAS,CME); R01HL151630 and NSF 2225554 (BJS), K99HL165072 (AMJ), R01HL164555 (SR)

