## supplement for "Siglec-F Deficiency Prevents Fibrosis After Bleomycin-Induced Acute Lung Injury"

### ONLINE DATA SUPPLEMENT

Marika Orlov[1], Sunad Rangarajan[1], Ana M. Jaramillo[1], Qihua Ye[1], Naoko Hara[1], Kenny Ngo[1], James C. NeeDell[1], Anna Q. Harder[1], Fan Jia[2], Brian Vestal[2], Rachel Z. Blumhagen[2], Ting-Hui Tu[3], Jazalle McClendon[3], Alexandra L. McCubbrey[1,3], Bradford J. Smith[4,5], David A. Schwartz[1,6], *William J. Janssen[1,3], and *Christopher M. Evans[1,6]

1. Division of Pulmonary Science and Critical Care Medicine, University of Colorado School of Medicine, Denver, CO, USA
2. Center for Genes, Environment and Health, National Jewish Health, Denver, CO
3. Department of Medicine, Division of Pulmonary, Critical Care and Sleep Medicine, National Jewish Health, Denver, CO, USA
4. Department of Bioengineering, University of Colorado Denver | Anschutz Medical Campus, Aurora, CO, USA
5. Section of Pulmonary and Sleep Medicine, Department of Pediatrics, University of Colorado Denver, Anschutz Medical Campus, Aurora, CO 80045, USA
6. Research Service, Rocky Mountain Regional Veterans Affairs Medical Center, Aurora, Colorado, USA.

* These authors contributed equally to this work

**Methods**

Mice: All studies were conducted with the approval of the Institutional Animal Care and Use Committees at the University of Colorado and National Jewish Health. C57BL/6J mice were purchased from the Jackson Laboratories (Jax, Bar Harbor, ME). Housing rooms were maintained at 22°C, 30-40% humidity, and a 14/10 (h/h) light dark cycle. *Siglecf* knockout (KO, -/-) mice were provided by Dr. Ajit Varki (University of California San Diego) on a congenic C57BL/6J background and maintained by breeding with C57BL/6J. Wild type (WT, +/+) littermates and purchased C57BL/6J mice were used as noted. Due to sexual dimorphism with the bleomycin model, 8-12 week old male mice were used in these studies (1).

Murine Bleomycin Injury: A bleomycin challenge model of acute lung injury and fibrosis was applied in accordance American Thoracic Society guidelines (2). Bleomycin (1.5-2.5 units/kg, Meitheal Pharmaceuticals) or PBS was administered intratracheally (IT) to isoflurane-sedated mice using a modified gavage needle (1). On days 7, 14, and 21 after challenge, mice were sedated with urethane (2.0 g/kg, i.p.), euthanized by exsanguination under anesthesia, and endpoints were collected as described below.

Murine Bronchoalveolar Lavage: After euthanasia, the trachea was exposed, and an 18-gauge Luer stub was inserted into an incision made on the ventral portion and secured with silk thread.

For studies including histologic and other endpoints, a plug was placed into the opening of the tracheal cannula, and the chest wall was opened. The left lung was clamped at the mainstem bronchus, and the right lung was lavaged by instilling and removing 0.5 ml PBS containing 0.6 mM EDTA three times (PBS/EDTA, 1.5 ml total). BAL fluid was used for hemocytometer counts, differentials, and flow cytometry as reported previously (3, 4). Following lavage, the right lung was excised, snap frozen, and stored at -80° C for use in hydroxyproline assays, and the left lung was collected for histology as described below.

For RNAseq studies, BAL was performed as above except that 1 ml PBS/EDTA was used to lavage both lungs five times (5 ml total). BAL fluid was centrifuged at 300 x g, and cells were used for macrophage sorting and RNA isolation as described below.

Murine Histology: The left lung was held at functional residual capacity, ligated with silk thread, excised, and placed in methacarn solution for 24 hours at 4°C. Subsequently, the lung was cut into 5-8 sections, processed in paraffin, sectioned at 5 µm thickness, and collected on glass slides. For collagen staining, slides were deparaffinized, serially hydrated, and stained for 1 h in pico-sirius red solution (1% w/v Direct Red 80 (Sigma, St. Louis, MO), in a saturated aqueous solution of picric acid, Fisher Scientific). Slides were then washed twice in acidified water (0.087 N acetic acid), rinsed in tap water, serially dehydrated in graded ethanol and xylene, and mounted with Cytoseal.

Murine Hydroxyproline Assay (HP): HP assay was performed as previously described (5). Briefly, after lavage for BAL fluid, the right lung was isolated and weighed. The lung was then placed in a collection tube and PBS was added such that the final weight (lung + PBS) equilibrated to 1 ml to standardize the amount of lung tissue added to the HP reaction. Lungs were thawed, homogenized using an MPBio Fast Prep 24 grinder and lysis system, transferred to glass vials (Scientific resources #78315), and mixed with 12 N HCl acid in a 1:1 ratio of homogenized lung:acid. Tubes were then sealed and placed in a 100° C heat block overnight. The following morning tubes were removed, cooled to room temperature, and 5 µl of lung:acid solution was mixed with 100 µl of 0.06 M chloramine T in citrate-acetate buffer, pH 6, in 96-well

ELISA, flat bottom plates. Samples were run in duplicate. Standards were made using cis-4-hydroxy-1 proline (Sigma) with a concentration range of 400 µg/ml to 03.9 µg/ml and were also run in duplicate. Plates were incubated at room temperature for 20 min, 100 µl of Ehrlich's solution (1.2 M dimethylaminobenzaldehyde in 22% perchloric acid-*n*-propanol) was added to each well and incubated at 65°C for 20 min. Plates were read at 550 nm absorbance (Synergy HTX plate reader, Biotek, Winooski, VT) and concentrations were calculated using a standard curve.

<u>Murine Stereologic Collagen Quantification:</u> Sirius red stained slides were imaged at 200x total magnification on an Olympus VS120 upright scanning microscope under brightfield and polarized illumination (Figure E1). Stereologic quantification was performed in accordance with American Thoracic Society guidelines (6). Images of whole lung sections were pre-processed to remove non-lung image data (white space outside of lung tissues). Images were then segmented into 533 x 400 µm sub-samples by standardized uniform random sampling (~40-75 images per mouse) that were imported into a custom application for quantitative analysis developed in Matlab (Mathworks, Natick, MA).

Each subsample contained three image layers: a polarized image, a brightfield view of the same area, and a grid overlay. Grids consisted of 48 uniformly distributed boxes, which were used to manually assign tissue compartments using the brightfield image. The following tissue categories were used: bronchial airway, bronchial vessel, airspace, and parenchyma. Parenchyma was defined as bronchioles and small vessels lacking smooth muscle, alveolar septa, alveolar airspace, alveolar duct airspace, and pleura.

To limit bias, assignments to tissue categories were made using brightfield images without knowledge of whether underlying polarized pixels were detected. Studies were also performed by two blinded investigators who did not have knowledge of mouse treatment or genotype until after completion.

For fibrosis quantification, the volume fraction of parenchymal collagen and distribution of collagen signals were measured. Volume fractions were calculated as the numbers (N) of positive collagen (col) signals in parenchyma (par) under polarized light relative total number of points identifying parenchymal (par) tissue and air. The distribution of collagen was quantified by determining the numbers of collagen signals that were adjacent to other collagen signals (col,adj) relative to total parenchymal collagen (col,par) in lung images. These measurements are shown in Equations 1 and 2, where ‘i’ is images that number from 1 to ‘n’ for each mouse. Data were computed as numbers of points per mouse (not averages of fractions per image).

**<u>Equation 1.</u>** Parenchymal collagen volume fraction.

$$Vv = \sum_{i=1}^{n} N_{(col,par)} \div \sum_{i=1}^{n} N_{(\mathrm{par})}$$

**<u>Equation 2.</u>** Parenchymal collagen adjacency.

$$Adj = \sum_{i=1}^{n} N_{(col,adj)} \div \sum_{i=1}^{n} N_{(col,par)}$$

Stereology results were validated by comparing with Ashcroft and hydroxyproline assay results (Figure E2) (2).

<u>Murine Modified Ashcroft Scoring:</u> Two independent and blinded researchers performed histological analysis of 30-40 Sirius red sub-sampled images per mouse (obtained as described above). Each image was given a score of 0-8 based on the grade of fibrosis as described in the modified Ashcroft Scale by Hubner, et al (7). Scores were averaged per mouse, per observer.

All mice that had differences in mean scores equal to or greater than 1 were counted by a third blinded researcher.

Murine Macrophage Sorting: BAL fluid was collected as above and centrifuged at 300 x g for 5 min at 4°C and then labeled with the following markers using an antibody dilution of 1:200 in the presence of FcBlock: CD45 FITC (Clone 30-F11, BD), CD64 PE-Cy7 (Clone X54-5/7.1, Biolegend), CD88 APC (Clone 20/70, Biolegend), Ly6G Pacific Blue (Clone 1A8, Biolegend) at 4°C for 45 min. The cells were then centrifuged at 300 x g for 5 min, washed in FACS buffer, and re-suspended in 500 µl FACS buffer. Prior to sorting, 1 drop of NucBlue (Invitrogen) was added to the cells. Alveolar macrophages were gated as CD45+CD64+CD88+Ly6G- cells on a Sony Biotechnology SY3200 sorter.

Murine RNA seq: RNA was isolated from sorted cells using Qiagen micro RNA kits. RNA purity, quantity, and integrity were determined by NanoDrop (ThermoFisher Scientific, Waltham, MA) and TapeStation 4200 (Agilent, Santa Clara, CA) analyses prior to RNA-seq library preparation. The Universal Plus mRNA-Seq library preparation kit with NuQuant (Tecan, Männedorf, Switzerland) was used with an input of 200 ng of total RNA to generate RNA-Seq libraries. Paired-end sequencing reads of 150 bp was generated on NovaSeq 6000 (Illumina, La Jolla, CA) sequencer at a target depth of 80 million paired-end reads per sample. Raw sequencing reads were de-multiplexed using bcl2fastq and processed using the nf-core/rnaseq pipeline (3.12.0) with nextflow (23.04.1). Resulting count data were transformed using variance stabilizing transformation (VST) (DESeq2, 1.42.0), and differentially expressed genes (DEGS) were identified using limma (3.58.1) in R (4.3.1).

Human BAL: Alveolar macrophages were isolated from freshly explanted human donor lungs as previously described (8, 9). Donors were non-smokers that died from non-pulmonary causes, had clear chest radiographs and PaO2:FiO2 ratios > 300. Right middle lobes were lavaged with 500 ml room temperature PBS. Fluid was passed through a 100 μl strainer centrifuged at 500 x g for 15 minutes. Freshly isolated cells were used for flow cytometry.

Human Macrophage Flow Cytometry. BAL was collected as above and centrifuged at 300 x g for 5 min at 4°C and then labeled with the following markers in the presence of FcBlock: HLA-DR eFluor450 (Clone L243, eBioscience), CD116 FITC (Clone 4H1, Biolegend), CD16 PerCP-Cy5.5 (Clone CB16, eBioscience), CD14 PE-Cy7 (Clone MφP9, BD BioSciences), Siglec 7 (polyclonal Rabbit IgG, unconjugated, Invitrogen), Siglec 8 (polyclonal Rabbit IgG, unconjugated, Sigma), Siglec 9 PE (Clone K8, Biolegend) at 4°C for 45 min. The cells were then spun down at 300 x g for 5 min and washed in FACS buffer. Siglec 7 and Siglec 8 primary antibodies and their isotype controls were stained with Cy3 $F(ab')_2$ fragment donkey anti-rabbit IgG (Jackson Immuno Research Labs) for 20 min on ice. Isotype control for Siglec 9 was PE-conjugated mouse IgG1. The cells were then centrifuged at 300 x g for 5min, washed in FACS buffer, and re-suspended in 500 µl FACS buffer. Prior to sorting, 1 drop of NucBlue (Invitrogen) was added to the cells. Cells were analyzed on an LSR II flow cytometer (BD Biosciences). Cells were identified using forward-scatter and side-scatter, and doublets and dead cells were excluded. Macrophages were identified by high side-scatter and positive staining for CD14 and HLA-DR. Siglec expression was compared to both fluorescence minus one (FMO) and isotype control.

Human Tissue Immunofluorescence: Lung tissue specimens from patients with idiopathic pulmonary fibrosis (IPF) were obtained from the NHLBI Lung Tissue Research Consortium (LTRC; https://ltrcpublic.com/). Studies were performed in compliance with local Institutional Review Boards, and individuals gave written informed consent to participate.

Slides containing human IPF lung tissue were de-paraffined in Histochoice, rehydrated, and then immersed in 1x target retrieval solution (Agilent) at high pressure for 20 min in an InstaPot, and washed in Tween-Tris-buffered saline (TTBS) twice. The slides were then washed in 0.2% triton X-100 in TTBS for 10 min, then washed in TTBS twice and blocked in 5% donkey and 5% goat serum in TTBS for 1 hour at room temperature. Primary antibodies were added that included either CD68-PE (Clone Y1/82A, Biolegend, 1:50), Siglec-7 (Clone EPR26252-79, Abcam, dilution 1:200) or CD68-PE, Siglec-9 (OTI1D9, Novus, dilution 1:50), or Isotype controls (Mouse IgG2bk-PE, Clone MPC-11, Biolegend; Polyclonal Rabit IgG, Abcam; Mouse IgG1k, MOPC-21, BD). Target proteins were labeled overnight at 4˚C. Slides were washed twice in TTBS and then incubated with Alexa fluor (AF) 750-conjugated goat anti-mouse for Siglec-9 and AF647-conjugated donkey anti-rabbit AF647 for Siglec 7) secondary antibodies (1:200, 1 hour, room temperature). Slides were then washed in TTBS twice and stained with Tomato lectin-FITC (Vector, dilution 1:300) for 30 min at room temperature. All tissue sections were treated with Autoquencher (Vector) to reduce autofluorescence before mounting the slides. Slides were imaged on an Olympus VS200.

Statistical Analysis: Statistical analyses were conducted in Prism 10 (GraphPad, La Jolla, CA) as described in the text. Data were analyzed using t-Tests for two-sample comparisons with normal distributions. For skewed or non-normally distributed two-sample data, Mann-Whitney U tests were applied. Multiple comparisons were made using ANOVA or Kruskal-Wallis tests for normally and non-normally distributed data, respectively. Dunn's and Dunnet's post-hoc tests was used to compare specified pairs or differences from bleomycin challenged wild type, respectively. A p-value cut-off of 0.05 was applied.

For RNA-Seq data, genes were first filtered to keep only those that had a counts per million value > 5 in at least 3 samples (i.e. the size of the smallest experimental group of interest). Subsequently, the digital counts were transformed into continuous measures using the variance stabilizing transformation implemented in the DESeq2 R package v1.38.3 (10). A Linear model using these transformed expression values was fit for each gene using a cell-means parameterization for groups defined by all pairwise combinations of genotype and days since challenge using the limma R package v3.54.2 (11). Linear contrasts were used to compute fold changes and p-values based upon moderated t-tests on these differences. Multiple comparisons adjustments were done using the Benjamini-Hochberg method for controlling false discovery rates (FDRs), and an FDR threshold of 0.05 was used to determine statistical significance (12). Gene lists for each comparison of interest were input in the enrichR R package v3.2 9 (https://cran.r-project.org/web/packages/enrichR/index.html) to look for overrepresentation using GO Biological Processes as the annotation database for pathways (13).

**Table E1**. Macrophages, lymphocytes and neutrophils in the BAL fluid 7, 14, and 21 days after bleomycin challenge in *Siglecf*$^{-/-}$ and WT mice. P-values depict results of two-way Mann-Whitney analyses.

| Day 7 Cells (x10$^4$/ml) | | *Siglecf* WT (mean +/- SEM) | *Siglecf* KO (mean +/- SEM) | Mann-Whitney test *p-value* |
|---|---|---|---|---|
| | | N=7 | N=5 | |
| | Macrophages | 21.5 +/- 4.2 | 18.6 +/- 7.9 | *0.32* |
| | Lymphocytes | 2.8 +/- 1 | 0.8 +/- 0.4 | *0.11* |
| | Neutrophils | 1.5 +/- 0.6 | 0.9 +/- 0.5 | *0.53* |
| Day 14 (x10$^4$/ml) | | | | |
| | | N=6 | N=10 | |
| | Macrophages | 31.5 +/- 10.6 | 26.3 +/- 5.0 | *0.87* |
| | Lymphocytes | 2.4 +/- 1.4 | 2.2 +/- 0.6 | *0.49* |
| | Neutrophils | 0.5 +/- 0.3 | 0.4 +/- 0.2 | *0.94* |
| Day 21 (x10$^4$/ml) | | | | |
| | | N=13 | N=20 | |
| | Macrophages | 35.9 +/- 4.3 | 28 +/- 2.5 | *0.12* |
| | Lymphocytes | 5 +/- 0.9 | 5 +/- 1 | *0.65* |
| | Neutrophils | 2.8 +/- 0.9 | 0.4 +/- 0.1 | *0.01* |

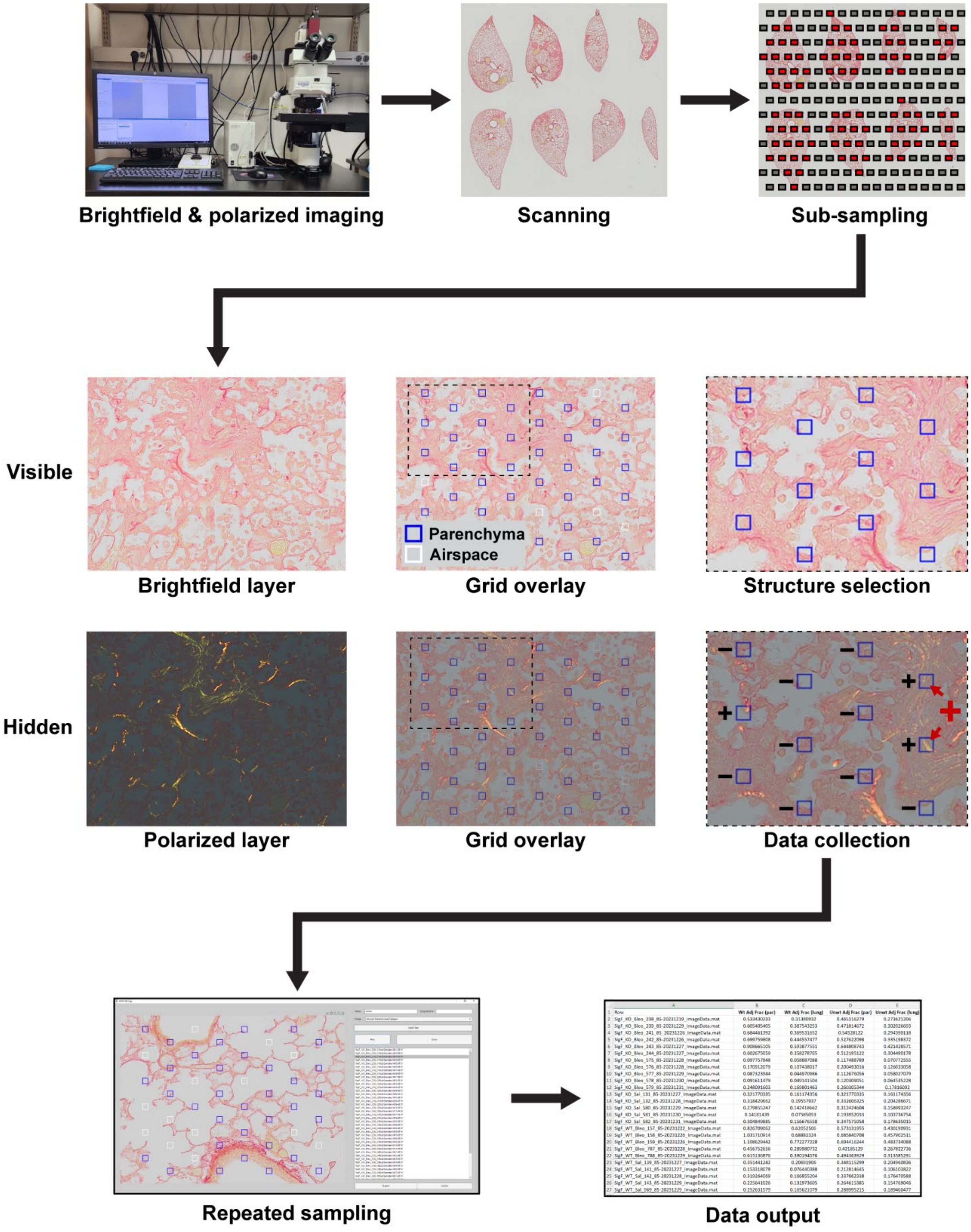


**Figure E1. Imaging and stereology workflow.** In Data collection panel, black '**+**' or '**−**' indicates positive or negative for collagen. Red '**+**' and arrows indicate adjacent collagen positive signals.

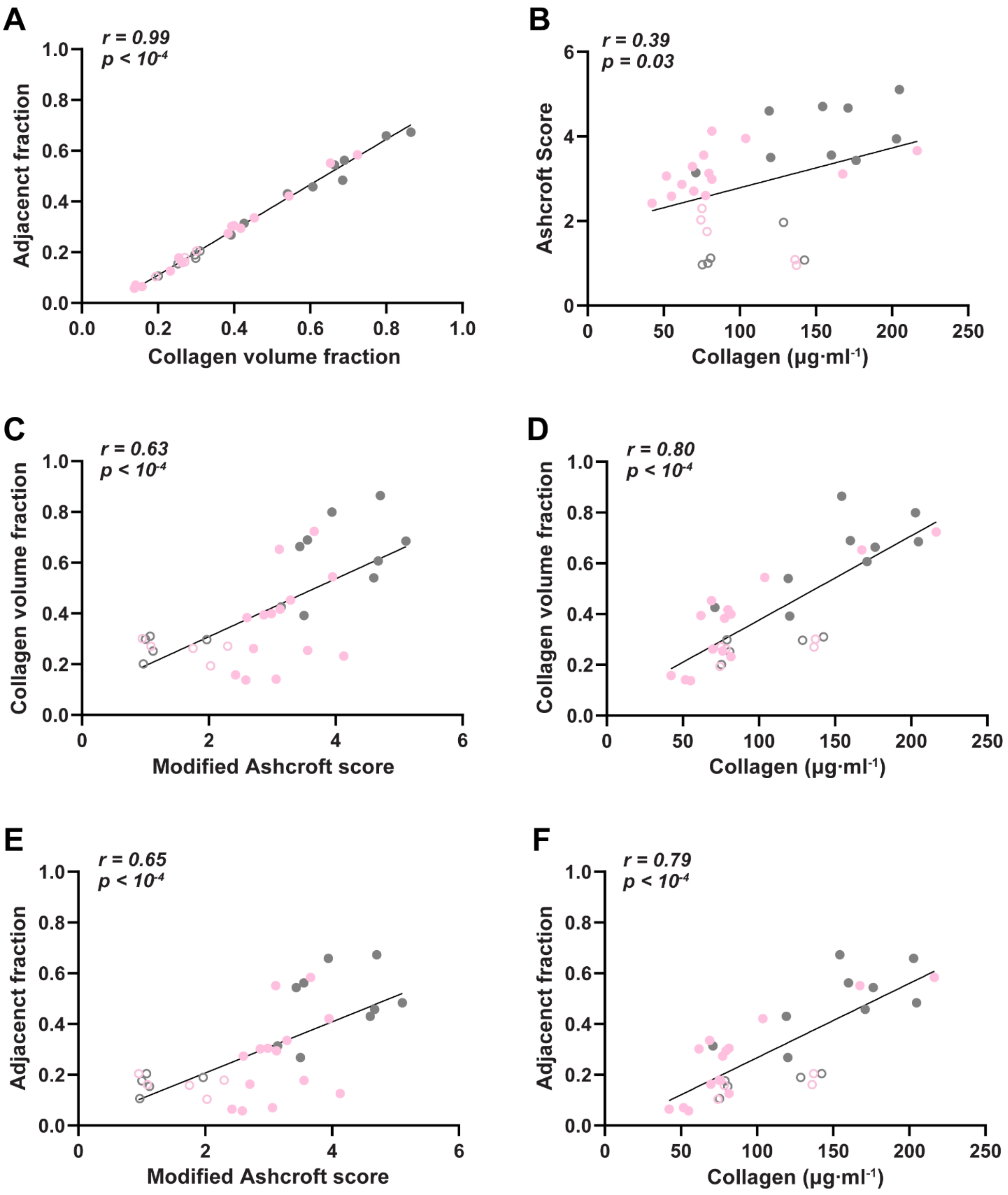


**Figure E2. Stereology validation. A.** Adjacency of signals used to estimate collagen density were compared to overall collagen volume fractions. **B-F.** Correlations were assessed among collagen volume density, adjacency, Ashcroft scoring, and hydroxyproline assay results. Ashcroft scoring and hydroxyproline content (collagen concentration) showed weak, but statistically significant correlations (**B**) that were improved by comparisons with collagen volume fraction (**C,D**) and adjacency (**E,F**). Spearman correlations (r) and p-values are shown.

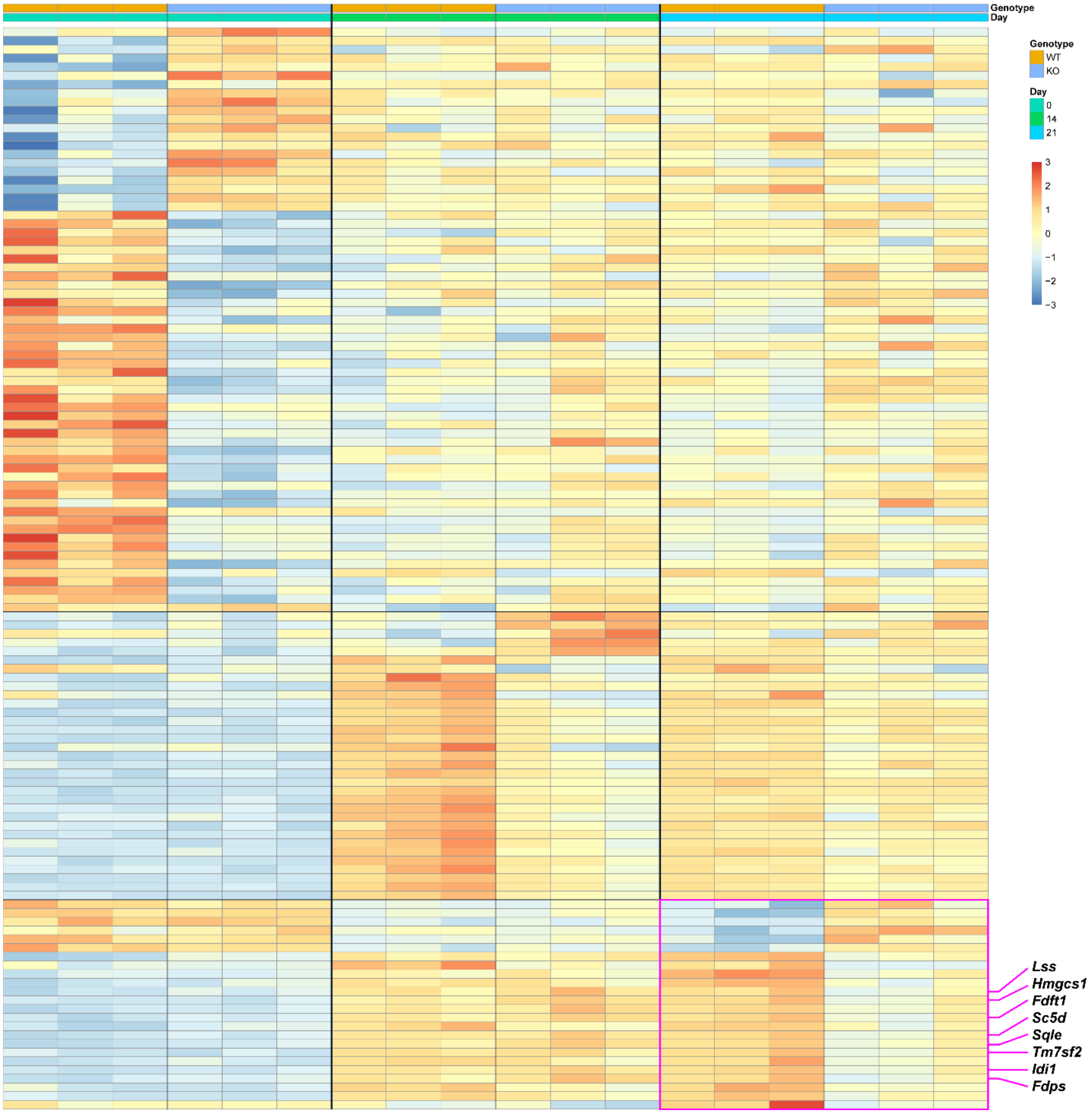


**Figure E3. Heatmap of bulk RNAseq analysis.** Airspace macrophage gene expression from BAL of naive (day 0) and bleomycin challenged (days 14 and 21) was compared between WT and *Siglecf* KO mice. Each column represents an individual mouse, and each row represents a single gene. There is decreased expression of genes involved in lipid and cholesterol metabolism at day 21 (magenta box). Transcripts identified at lower right are displayed in Figure 1I.
